# An experimentally informed evolutionary model improves phylogenetic fit to divergent lactamase homologs

**DOI:** 10.1101/003848

**Authors:** Jesse D. Bloom

## Abstract

Phylogenetic analyses of molecular data require a quantitative model for how sequences evolve. Traditionally, the details of the site-specific selection that governs sequence evolution are not known *a priori*, making it challenging to create evolutionary models that adequately capture the heterogeneity of selection at different sites. However, recent advances in high-throughput experiments have made it possible to quantify the effects of all single mutations on gene function. I have previously shown that such high-throughput experiments can be combined with knowledge of underlying mutation rates to create a parameter-free evolutionary model that describes the phylogeny of influenza nucleoprotein far better than commonly used existing models. Here I extend this work by showing that published experimental data on TEM-1 beta-lactamase (Firnberg et al., 2014) can be combined with a few mutation rate parameters to create an evolutionary model that describes beta-lactamase phylogenies much than most common existing models. This experimentally informed evolutionary model is superior even for homologs that are substantially diverged (about 35% divergence at the protein level) from the TEM-1 parent that was the subject of the experimental study. These results suggest that experimental measurements can inform phylogenetic evolutionary models that are applicable to homologs that span a substantial range of sequence divergence.

## Introduction

Most approaches for the phylogenetic analysis of gene sequences require a quantitative evolutionary model specifying the rate at which each site substitutes from one identity to another. In maximum-likelihood and Bayesian approaches, the evolutionary model is used to calculate the likelihood of the observed sequences given the phylogenetic tree (Felsenstein, 1981; Huelsenbeck et al., 2001). In distance-based approaches, the evolutionary model is used to calculate the distances between pairs of sequences (Saitou and Nei, 1987; Hasegawa et al., 1985). For all these approaches, inaccurate evolutionary models can lead to errors in inferred phylogenetic properties, including incorrect estimates of evolutionary distances (Halpern and Bruno, 1998) and incorrect tree topologies (Felsenstein, 1978; Huelsenbeck and Hillis, 1993; Lartillot et al., 2007; Rokas and Carroll, 2008).

Unfortunately, existing phylogenetic evolutionary models are extreme simplifications of the actual process of mutation and selection that shapes sequence evolution (Thorne et al., 2007). At least two major unrealistic assumptions afflict most of these models. First, in order to make phylogenetic algorithms computationally tractable, it is generally assumed that each site within a gene evolves independently. Second, most widely used evolutionary models compound the first assumption of independence among sites with the second unrealistic assumption that all sites evolve identically – a severely flawed assumption since there is overwhelming evidence that proteins have strong preferences for certain amino acids at specific sites (Ashenberg et al., 2013; Halpern and Bruno, 1998). It is the second of these unrealistic assumptions that is addressed by the experimentally informed evolutionary model described here.

A major reason that most phylogenetic evolutionary models assume that sites evolve identically is because it has traditionally been thought that there is insufficient information to do better. In the absence of *a priori* knowledge about selection on individual sites, the parameters of an evolutionary model must be inferred from the same sequences that are being analyzed phylogenetically. For instance, typical codon-level models infer parameters describing the equilibrium frequencies of different codons, the relative rates of transition and transversion mutations, the relative rates of nonsynonymous and synonymous mutations, and in many cases the shapes of distributions that allow some of these rates to be drawn from several categories (Goldman and Yang, 1994; Muse and Gaut, 1994; Yang et al., 2000; Kosiol et al., 2007). These parameters can easily be inferred for a single general model that applies to all sites in a gene, but it is much more challenging to infer them separately for each site without overfitting the available sequence data (Posada and Buckley, 2004; Rodrigue, 2013). Some studies have attempted to bypass this problem by predicting site-specific substitution rates or classifying sites based on knowledge of the protein structure (Thorne et al., 1996; Goldman et al., 1998; Scherrer et al., 2012; Rodrigue et al., 2009; Kleinman et al., 2010) – however, such approaches are limited by the fact that the relationship between protein structure and site-specific selection is complex, and cannot be reliably predicted even by state-of-the-art molecular modeling (Potapov et al., 2009).

A more recent and more promising alternative approach is to infer the site-specific substitution process directly from the sequence data (Lartillot and Philippe, 2004; Rodrigue et al., 2010; Le et al., 2008; Wu et al., 2013; Wang et al., 2008). Fully specifying a different substitution process for each site in a maximum-likelihood framework requires inferring a large number of free parameters (there are 19 *× L* parameters for a gene with *L* codons if selection is assumed to act on the amino-acid sequence). Therefore, the details of the site-specific substitution process must be inferred without overfitting the finite available data. Two strategies have been successfully employed to do this: constraining sites to fall in a fixed number of different substitution-model classes (Wang et al., 2008; Le et al., 2008), or using non-parametric Bayesian mixture models that treat the site-specific substitution probabilities as random variables that are integrated over a statistical distribution estimated from the data (Lartillot and Philippe, 2004; Rodrigue et al., 2010). Although both of these strategies are somewhat complex, they yield much more nuanced evolutionary models that eliminate some of the problems associated with the unrealistic assumption that sites evolve identically.

Even more recently, a new type of high-throughput experiment has begun to yield data that enables the creation of site-specific evolutionary models without any need to infer site-specific selection from the naturally occurring gene sequences that are the subject of the phylogenetic analysis. This new type of experiment is deep mutational scanning (Fowler et al., 2010; Araya and Fowler, 2011), a technique in which a gene is randomly mutagenized and subjected to functional selection in the laboratory, and then deep sequenced to quantify the relative frequencies of mutations before and after selection. In cases where the laboratory selection is sufficiently representative of the gene’s real biological function, these experiments provide information that can be used to approximate the site-specific natural selection on mutations. To date, deep mutational scanning has been used to quantify the impact of most nucleotide or codon mutations to several proteins or protein domains (Fowler et al., 2010; Roscoe et al., 2013; Starita et al., 2013; Melamed et al., 2013; Traxlmayr et al., 2012; McLaughlin Jr et al., 2012; Firnberg et al., 2014; Bloom, 2014). For a few of these studies, the experimental coverage of possible mutations is sufficiently comprehensive to define site-specific amino-acid preferences for all positions in a gene.

I have previously shown that such experimentally determined site-specific amino-acid preferences can be combined with measurements of mutation rates to create a parameter-free evolutionary model that describes the phylogeny of influenza nucleoprotein far better than existing non-site-specific models that contain numerous free parameters (Bloom, 2014). Here I extend that work by showing that it is also possible to create an experimentally informed evolutionary model for another gene. I do this using deep mutational scanning data published by Firnberg et al. (2014) that quantifies the effects of nearly all amino-acid mutations on TEM-1 beta-lactamase. In this case, no measurements of mutation rates are available, so I construct an evolutionary model that is informed by the experimentally measured site-specific amino-acid preferences but also contains a few free parameters representing the mutation rates. I also augment this model with an additional parameter that reflects the stringency of the site-specific amino-acid preferences in natural evolution versus the deep mutational scanning experiment used to measure these preferences. I show that this evolutionary model greatly improves the phylogenetic fit to both TEM and SHV beta-lactamases, the latter of which are substantially diverged (about 35% divergence at the protein level) from the TEM-1 parent that was the subject of the deep mutational scanning by Firnberg et al. (2014). These results generalize previous work on experimentally determined evolutionary models, and suggest that site-specific amino-acid preferences are sufficiently conserved during evolution to be applicable to gene homologs that span a substantial range of sequence divergence.

## Results

### Evolutionary model

#### Summary of evolutionary model

I have previously described a codon-level phylogenetic evolutionary model for influenza nucleoprotein for which both the site-specific amino-acid preferences and the nucleotide mutation rates (assumed to be identical across sites) were determined experimentally (Bloom, 2014). The current work examines a protein for which the site-specific amino-acid preferences have been measured experimentally, but for which the nucleotide mutation rates are unknown. It is therefore necessary to extend the evolutionary model to treat the nucleotide mutation rates as unknown free parameters. Here I describe this extension.

In the model used here, the rate *P_r,xy_* of substitution from codon *x* to some other codon *y* at site *r* is

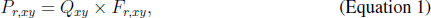

where *Q_xy_* denotes the rate of mutation from *x* to *y*, and *F_r,xy_* gives the probability that a mutation from *x* to *y* fixes if it occurs. This equation assumes that mutation rates are uniform across sites, and that the selection on mutations is site-specific but site-independent (i.e. the fixation probability at one site is not influenced by mutations at other sites).

#### Fixation probabilities from amino-acid preferences

The fixation probability of a mutation from codon *x* to *y* is assumed to depend only on the encoded amino acids 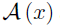 and 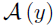, as synonymous mutations are assumed to be selectively neutral. The fixation probabilities *F_r,xy_* are defined in terms of the experimentally measured amino-acid preferences at site *r*, where *π_r,a_* denotes the preference for amino-acid *a* at site *r*, and the preferences at each site sum to one 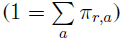.

As in previous work (Bloom, 2014), I consider two different mathematical relationships between the amino-acid preferences and the fixation probabilities. However, novel to this work, I consider a generalization of these relationships that allows the stringency of the amino-acid preferences to differ between natural sequence evolution and the deep mutational scanning experiments used to measure *π_r,a_*. Specifically, I take the probability of fixation to depend on (*π_r,a_*)*^β^* where *β* is a free parameter (constrained to have values ≥ 0) that scales the stringency of the amino-acid preferences. A value of *β* = 1 implies equally stringent preferences in natural evolution and the deep mutational scanning experiments. A value of *β <* 1 corresponds to less stringent preferences in natural evolution than in the deep mutational scanning experiments, while a value of *β >* 1 corresponds to more stringent preferences in natural evolution in the deep mutational scanning experiments. Naively, one might conjecture that *β* will typically have values *>* 1, since laboratory experiments tend to be less stringent than natural evolutionary selection. In the following sections, I will describe testing evolutionary models models that constrain *β* = 1 versus models that treat *β* as a free parameter.

With the addition of the stringency parameter *β*, I consider the following relationships between the amino-acid preferences and the fixation probabilities. The first relationship derives from considering the amino-acid preferences to be directly related to selection coefficients, and is a generalization of the ninth equation derived by Halpern and Bruno (1998):

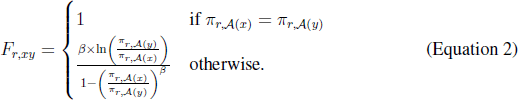

Note that in Equation 2 the mutation terms in the original equation derived by Halpern and Bruno (1998) are all set to be equal since the calculations of the amino-acid preferences *π_r,a_* from the deep mutational scanning experiments already correct for differences in the mutagenesis rates, whereas Halpern and Bruno (1998) were inferring the preferences from natural sequences and so had to account for the mutation rates during natural evolution. Note also that the ninth equation of Halpern and Bruno (1998) contains a typographical error in the denominator which is corrected in Equation 2. Halpern and Bruno (1998) derive Equation 2 with *β* = 1 by assuming that the sequences are evolving in the weak-mutation limit, and rigorous application of this relationship with *β* = 1 in the context of the current work requires assuming that the effective population size in the deep mutational scanning experiment is equivalent to that for the natural sequences that are being phylogenetically analyzed.

The second relationship is based on considering the amino-acid preferences to reflect the fraction of genetic backgrounds that tolerate specific mutations rather than selection coefficients in any one genetic background. Specifically, experiments have shown that mutations that are deleterious in one genetic background can sometimes be neutral (or even advantageous) in a related genetic background (Lunzer et al., 2010; Gong et al., 2013). One reason that mutational effects depend on genetic background is that most proteins are under selection to maintain their overall stability above some critical threshold (Gong et al., 2013). This type of threshold selection gives rise to the evolutionary dynamics described in Bloom et al. (2007), where stabilizing mutations are always tolerated but destabilizing mutations are only tolerated in a fraction of genetic backgrounds. In this case, mutations to higher preference (putatively more stabilizing) amino acids will always be able to fix without deleterious effect, but mutations to lower preference (putatively less stabilizing) amino acids are only sometimes tolerated. One way to represent these dynamics is to use a relationship equivalent to the Metropolis et al. (1953) sampling criterion:

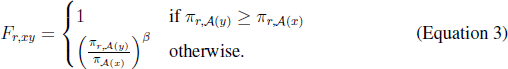

Both of these relationships (Equation 2 and Equation 3) share the feature that mutations to higher-preference amino acids fix more frequently than mutations to lower-preference amino acids as long as *β >* 0.

#### Mutation rates

The rate of mutation *Q_xy_* from codon *x* to *y* is defined in terms of the underlying rates of nucleotide mutation. Let *R_m→n_* denote the rate of mutation from nucleotide *m* to *n*. Then

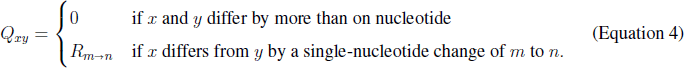

Assuming that the same mutation process operates on both the sequenced and complementary strands of the nucleic acid gives the constraint

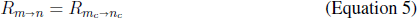

where *m_c_* denotes the complement of nucleotide *m*, since for example a mutation from *A* to *G* on one strand induces a mutation from *T* to *C* on the other strand. There are therefore six unique nucleotide mutation rates: *R_A→C_* = *R_T→G_*, *R_A→G_* = *R_T→C_*, *R_C→A_* = *R_T→A,_ R_C→A_* = *R_G→T_*, *R_C→G_* = *R_G→C_*, and *R_C→T_* = *R_G→A_*.

In principle, these six mutation rates could be measured experimentally for the system of interest. In my previous work on influenza nucleoprotein (Bloom, 2014), I was able to devise an experimental system for measuring the mutation rates for influenza in cell culture. The mutation rates measured for influenza in this system were roughly symmetric (*R_m→n_* = *R_n→m_*), which was sufficient to make the overall evolutionary model in Equation 1 reversible. However, in general it is unlikely to be feasible to measure the mutation rates for most systems of interest. Furthermore, it is known that mutation process is *AT*-biased for many species (Hershberg and Petrov, 2010), meaning that in general mutation rates will not be symmetric. Therefore, in general (and also for lactamase specifically) it is necessary to infer the mutation rates from the sequence data without assuming that they are symmetric.

In the absence of any constraints on the six mutation rates given above, the overall evolutionary model defined by Equation 1 will not necessarily be reversible. However, it turns out (see Methods) that placing the following constraint on the mutation rates is sufficient to make the overall evolutionary model reversible:

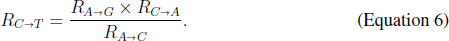

This constraint lacks a biological justification, and is assumed purely for the mathematical convenience that it makes the model reversible. Although there is no biological reason to believe that Equation 6 actually holds for real evolution, it is possible to give interpretations about what assuming this equation implies about the mutational process. One interpretation is that the probability of mutating from *C* to *G* via an intermediate mutation to *T* is equal to the probability of mutating from *C* to *G* via an intermediate mutation to *A*, as Equation 6 implies that *R_C→T_ × R_T→G_* = *R_C→A_ ^×^ R_A→G_*. Another interpretation is that the *AT* bias is the same for transitions and transversions, as Equation 6 implies that *R_C→T_ /R_T→C_* = *R_C→A_/R_A→C_*.

In the absence of independent information to calibrate absolute values for the branch lengths or mutation rates, one of the rates is confounded with the branch-length scaling and so can be assigned an arbitrary value *>* 0 without affecting the tree or its likelihood. Here the choice is made to assign

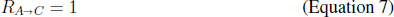

so that all other mutation rates are defined relative to this rate. With these constraints, there are now four independent mutation rates that must be treated as unknown free parameters:

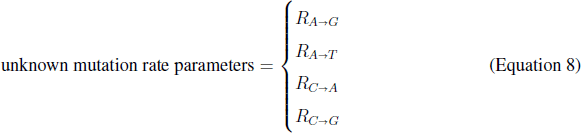

In practice, these four mutation rate parameters will be estimated at their maximum likelihood values given the sequences and tree topology.

#### Equilibrium frequencies

Calculation of a phylogenetic likelihood requires assigning evolutionary equilibrium frequencies to the possible codons in addition to specifying the transition probabilities given by Equation 1. In many conventional phylogenetic models, these equilibrium frequencies are treated as free parameters that are estimated empirically from the sequence data. However, in reality the equilibrium frequencies are the result of mutation and selection, and so can be calculated as the stationary state of the stochastic process defined by the evolutionary model. Specifically, it can be shown (see Methods) that for the evolutionary model in Equation 1, the equilibrium frequency *p_r,x_* of codon *x* at site *r* is

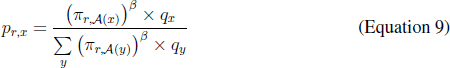

where *q_x_* is given by

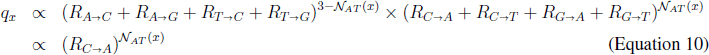

where 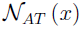 is the number of *A* and *T* nucleotides in codon *x*, the proportionality constant is never needed since *q_x_* values always appear in the form of ratios, and the simplification from the first to second line follows f rom Equation 5, Equation 6, and Equation 7. The equilibrium frequencies *p_r,x_* are therefore completely determined by knowledge of the experimentally determined amino-acid preferences *π_r,a_*, the mutation rate parameters in Equation 8, and the value of the stringency parameter *β*.

### Experimentally determined amino-acid preferences for beta-lactamase

The site-specific amino-acid preferences for beta-lactamase were determined using data from a previously published deep mutational scanning experiment performed by Firnberg et al. (2014). Specifically, Firnberg et al. (2014) created nearly all possible amino-acid mutants of TEM-1 beta-lactamase and then used antibiotic selection to enrich for functional variants at various antibiotic concentrations. Next, they used high-throughput sequencing to examine how the frequencies of mutations changed during this functional selection. They analyzed their data to estimate the impact of individual mutations on TEM-1 function, and had sufficient data to estimate the impact of 96% of the 297 *×* 19 = 5, 453 possible amino-acid mutations.

Firnberg et al. (2014) report the impact of mutations in terms of what they refer to as the “fitness” effects. Firnberg et al. (2014) calculate these fitness values from their deep mutational scanning experiment in such a way that a mutation’s fitness effect is approximately proportional to the highest concentration of antibiotic on which bacteria carrying that beta-lactamase variant are able to grow. Therefore, although the fitness values are not calculated in a true population-genetics framework, they certainly reflect effects of specific mutations on the ability of TEM-1 mutants to function.

Here I use the fitness values provided by Firnberg et al. (2014) to estimate the preferences for each of the 20 amino acids at each site in TEM-1. Specifically, let *w_r,a_* be the fitness value for mutation to amino-acid *a* at site *r* reported by Firnberg et al. (2014) in Data S2 of their paper. I calculate the preference *π_r,a_* for *a* at site *r* as

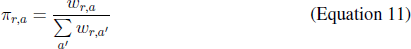

where the sum over *a*′ ranges over all 20 amino acids, the wild-type amino acid at site *r* is assigned a fitness of *w _r,a_* = 1 in accordance with the normalization scheme used by Firnberg et al. (2014), and the *w_r,a_* values for the 4% of mutations for which no value is estimated by Firnberg et al. (2014) are set to the average *w_r,a_* of all non-wildtype amino acids at site *r* for which a *w_r,a_* value is provided.

The amino-acid preferences calculated in this manner are displayed graphically in Figure 1 along with information about residue secondary structure and solvent accessibility (see Supplementary file 1 for numerical d ata). As is extensively discussed by Firnberg et al. (2014) in their original description of the data, these preferences are qualitatively consistent with known information about highly constrained positions in TEM-1, and show the expected qualitative patterns of higher preferences for specific (particularly hydrophobic) amino acids at residues that are buried in the protein’s folded structure. Here I focus on using these amino-acid preferences in a quantitative phylogenetic evolutionary model as described in the next section.

**Figure 1:**
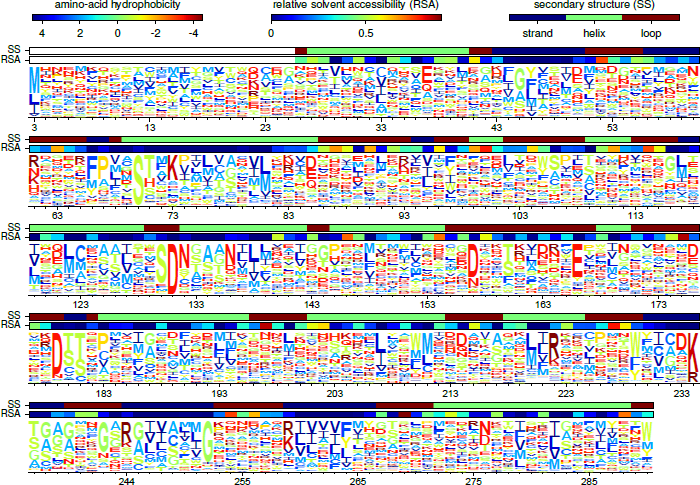
The amino-acid preferences for TEM-1 beta-lactamase, calculated from the data of Firnberg et al. (2014). The heights of letters are proportional to the preference for that amino acid at that position in the protein. Residues are numbered using the scheme of Ambler et al. (1991). Letters are colored according to the hydrophobicity of the amino acid. Bars above the letters indicate the secondary structure and relative solvent accessibility as calculated from the crystal structure in PDB entry 1XPB (Fonzé et al., 1995) using DSSP (Kabsch and Sander, 1983; Joosten et al., 2011), with maximum solvent accessibilities taken from Tien et al. (2013). The figure was generated using *WebLogo* (Crooks et al., 2004) integrated into the *mapmuts* software package (Bloom, 2014). The data and source code used to create this plot are provided via http://jbloom.github.io/phyloExpCM/example_2014Analysis_lactamase.html.

### Experimentally determined amino-acid preferences improve phylogenetic fit

#### TEM and SHV beta-lactamase phylogenetic trees

To test if evolutionary models informed by the experimentally determined amino-acid preferences are superior to existing alternative models, I compared the fit of various models to beta-lactamase sequence phylogenies. Firnberg et al. (2014) performed their deep mutational scanning on TEM-1 beta-lactamase. There are a large number of TEM beta-lactamases with high sequence identity to TEM-1; the next closest group of lactamases is the SHV beta-lactamases (Bush et al., 1995), which on average have 62% nucleotide and 65% protein identity to TEM beta-lactamases. I assembled a collection of TEM and SHV beta-lactamases from the manually curated Lahey Clinic database (http://www.lahey.org/Studies/). These sequences were aligned to TEM-1, and highly similar sequences (sequences that differed by less than four nucleotides) were removed. The resulting alignment contained 85 beta-lactamase sequences (Supplementary file 2), of which 49 were TEM and 36 were SHV.

Maximum-likelihood phylogenetic trees of the TEM and SHV beta-lactamases were constructed using *codonPhyML* (Gil et al., 2013) with the codon substitution model of either Goldman and Yang (1994) or Kosiol et al. (2007). The resulting trees are displayed in Figure 2. The two different substitution models give similar tree topologies – the Robinson-Foulds distance (Robinson and Foulds, 1981) between the trees inferred with the two different models is calculated by *RAxML* (Stamatakis, 2006) to be 0.14. In both cases, the trees partition into two clades of closely related sequences, corresponding to the TEM and SHV beta-lactamases.

**Figure 2:**
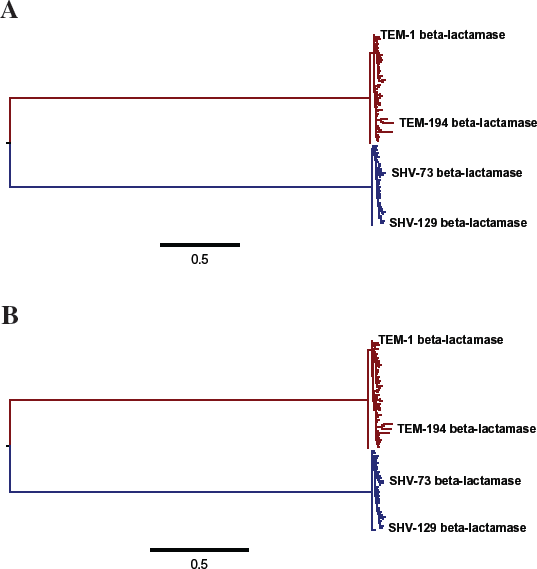
Phylogenetic trees of TEM (red) and SHV (blue) beta-lactamases inferred using *codon-PhyML* (Gil et al., 2013) with the codon substitution model of **(A)** Goldman and Yang (1994) or **(B)** Kosiol et al. (2007). The scale bars have units of number of codon substitutions per site. The inferred trees are similar for both models; the distance between the trees computed using the measure of Robinson and Foulds (1981) is 0.14. The TEM and SHV sequences each cluster into closely related clades: the average number of nucleotide and amino-acid differences between sequence pairs within these clades is 13 and 7 for the TEM sequences, and 10 and 5 for the SHV sequences. There is extensive divergence between these two clades: the average number of nucleotide and amino-acid differences between sequence pairs across the clades is 326 and 100. For both substitution models, a single transition-transversion ratio (*κ*) and four discrete gamma-distributed nonsynonymous-synonymous ratios (*ω*) were estimated by maximum likelihood. The equilibrium codon frequencies were determined empirically using the CF3x4 method (Pond et al., 2010) for the model of Goldman and Yang (1994), or the F method for the model of Kosiol et al. (2007) The data and source code used to create these trees are provided via http://jbloom.github.io/phyloExpCM/example_2014Analysis_lactamase.html.

#### Experimentally informed models are superior for combined TEM and SHV phylogeny

To compare the evolutionary models, *HYPHY* (Pond et al., 2005) was used to optimize the branch lengths and free parameters of the evolutionary models to their maximum likelihood values on the fixed tree topologies in Figure 2. This analysis showed that the evolutionary models informed by the experimentally determined amino-acid preferences were clearly superior to commonly used alternative codon-substitution models.

Specifically, Table 1 and Table 2 show that the experimentally informed evolutionary models fit the combined TEM and SHV phylogeny with higher likelihoods than any of a variety of commonly used alternative models, regardless of which tree topology from Figure 2 is used. This superiority is despite the fact that the alternative models (Goldman and Yang, 1994; Kosiol et al., 2007) contain many more free parameters. For instance, the most heavily parameterized alternative model contains 60 empirically estimated equilibrium frequency parameters plus an optimized parameter corresponding to the transition-transversion ratio, two optimized parameters corresponding to a gamma distribution of nonsynonymous-synonymous ratios across sites (Yang et al., 2000), and an optimized parameter corresponding to a distribution of substitution rates across sites (Yang, 1994). In contrast, the experimentally informed models only contain four or five free parameters (the mutation rates and optionally the stringency parameter *β*) – yet these experimentally informed models have substantially higher likelihoods. When AIC (Posada and Buckley, 2004) is used to penalize parameters, the superiority of the experimentally informed models is even more clear.

**Table 1:** Experimentally informed evolutionary models fit the combined TEM and SHV beta-lactamase phylogeny in Figure 2A much better than models that do not utilize experimental data. Shown are the difference in AIC relative to the best model (smaller ΔAIC indicates better fit), the log likelihood, the number of free parameters, and the values of key parameters. For each model, the branch lengths and model parameters were optimized for the fixed tree topology in Figure 2A. The “experimental” models use amino-acid preferences derived from the data of Firnberg et al. (2014) plus four mutation rate parameters (Equation 8) and optionally the stringency parameter *β*. For the “randomized” models, the experimentally measured amino-acid preferences are randomized among sites – these models are far worse since the preferences are no longer assigned to the correct positions. For the “avg. frequencies” models, the amino-acid preferences are identical across sites and are set to the average frequency of that amino acid in the entire lactamase sequence alignment – these models are also far worse than the experimentally informed models, since they do not utilize site-specific information. Fitting the stringency parameter to a value of *β >* 1 improves the fit of the experimentally informed models by enhancing the importance of the site-specific amino-acid preferences. Fitting the stringency parameter to a value of *β <* 1 improves the fit of the randomized and avg. frequencies model by effectively equalizing the preferences across amino acids. “GY94” denotes the model of Goldman and Yang (1994) with 9 equilibrium frequency parameters calculated using the CF3×4 method (Pond et al., 2010). “KOSI07+F” denotes the model of Kosiol et al. (2007) with 60 equilibrium frequency parameters calculated using the F methods. All variants of GY94 and KOSI07+F have a single transition-transversion ratio (*κ*) estimated by maximum likelihood. Different model variants either have a single nonsynonymous-synonymous ratio (*ω*) or values drawn from four discrete gamma-distributed categories (Yang et al., 2000), and either a single rate or rates drawn from four discrete gamma-distributed categories (Yang, 1994). The data and source code used to generate this table are provided via http://jbloom.github.io/phyloExpCM/example_2014Analysis_lactamase.html.

| model | ΔAIC | log<br>likelihood | parameters<br>(optimized +<br>empirical) | optimized parameters |
| --- | --- | --- | --- | --- |
| experimental, Equation 2, free $\beta$ | 0.0 | -4020.6 | 5 (5 + 0) | $R_{A \rightarrow G} = 2.3, R_{A \rightarrow T} = 0.6, R_{C \rightarrow A} = 0.8, R_{C \rightarrow G} = 0.7, \beta = 1.6$ |
| experimental, Equation 2, $\beta = 1$ | 46.4 | -4044.8 | 4 (4 + 0) | $R_{A \rightarrow G} = 2.3, R_{A \rightarrow T} = 0.6, R_{C \rightarrow A} = 0.8, R_{C \rightarrow G} = 0.8$ |
| experimental, Equation 3, free $\beta$ | 77.3 | -4059.3 | 5 (5 + 0) | $R_{A \rightarrow G} = 2.2, R_{A \rightarrow T} = 0.6, R_{C \rightarrow A} = 0.8, R_{C \rightarrow G} = 0.7, \beta = 1.2$ |
| experimental, Equation 3, $\beta = 1$ | 85.7 | -4064.5 | 4 (4 + 0) | $R_{A \rightarrow G} = 2.2, R_{A \rightarrow T} = 0.6, R_{C \rightarrow A} = 0.8, R_{C \rightarrow G} = 0.8$ |
| GY94, gamma $\omega$, gamma rates | 392.4 | -4208.8 | 13 (4 + 9) | $\kappa = 2.8, \omega \text{ shape} = 0.4, \text{mean } \omega = 0.7, \text{rate shape} = 1.2$ |
| KOSI07+F, gamma $\omega$, gamma rates | 410.7 | -4167.0 | 64 (4 + 60) | $\kappa = 0.3, \omega \text{ shape} = 0.4, \text{mean } \omega = 4.5, \text{rate shape} = 1.4$ |
| GY94, gamma $\omega$, one rate | 460.9 | -4244.1 | 12 (3 + 9) | $\kappa = 2.7, \omega \text{ shape} = 0.3, \text{mean } \omega = 1.0$ |
| KOSI07+F, gamma $\omega$, one rate | 467.0 | -4196.1 | 63 (3 + 60) | $\kappa = 0.3, \omega \text{ shape} = 0.3, \text{mean } \omega = 5.3$ |
| GY94, one $\omega$, gamma rates | 528.9 | -4278.1 | 12 (3 + 9) | $\kappa = 2.5, \omega = 0.4, \text{rate shape} = 1.2$ |
| KOSI07+F, one $\omega$, gamma rates | 551.3 | -4238.3 | 63 (3 + 60) | $\kappa = 0.4, \omega = 2.1, \text{rate shape} = 1.5$ |
| KOSI07+F, one $\omega$, one rate | 632.9 | -4280.1 | 62 (2 + 60) | $\kappa = 0.4, \omega = 2.0$ |
| GY94, one $\omega$, one rate | 656.2 | -4342.7 | 11 (2 + 9) | $\kappa = 2.4, \omega = 0.4$ |
| randomized, Equation 3, free $\beta$ | 724.5 | -4382.9 | 5 (5 + 0) | $R_{A \rightarrow G} = 2.4, R_{A \rightarrow T} = 0.6, R_{C \rightarrow A} = 0.9, R_{C \rightarrow G} = 0.9, \beta = 0.1$ |
| randomized, Equation 2, free $\beta$ | 735.1 | -4388.2 | 5 (5 + 0) | $R_{A \rightarrow G} = 2.4, R_{A \rightarrow T} = 0.6, R_{C \rightarrow A} = 0.9, R_{C \rightarrow G} = 0.9, \beta = 0.0$ |
| avg. frequencies, Equation 3, free $\beta$ | 820.8 | -4371.0 | 65 (5 + 60) | $R_{A \rightarrow G} = 2.4, R_{A \rightarrow T} = 0.6, R_{C \rightarrow A} = 1.0, R_{C \rightarrow G} = 0.9, \beta = 0.5$ |
| avg. frequencies, Equation 2, free $\beta$ | 841.8 | -4381.5 | 65 (5 + 60) | $R_{A \rightarrow G} = 2.4, R_{A \rightarrow T} = 0.6, R_{C \rightarrow A} = 1.0, R_{C \rightarrow G} = 0.9, \beta = 0.3$ |
| avg. frequencies, Equation 3, $\beta = 1$ | 858.0 | -4390.6 | 64 (4 + 60) | $R_{A \rightarrow G} = 2.3, R_{A \rightarrow T} = 0.6, R_{C \rightarrow A} = 1.1, R_{C \rightarrow G} = 1.0$ |
| avg. frequencies, Equation 2, $\beta = 1$ | 900.7 | -4412.0 | 64 (4 + 60) | $R_{A \rightarrow G} = 2.4, R_{A \rightarrow T} = 0.6, R_{C \rightarrow A} = 1.1, R_{C \rightarrow G} = 1.0$ |
| randomized, Equation 3 | 1264.9 | -4654.1 | 4 (4 + 0) | $R_{A \rightarrow G} = 2.3, R_{A \rightarrow T} = 0.6, R_{C \rightarrow A} = 0.9, R_{C \rightarrow G} = 0.9$ |
| randomized, Equation 2 | 1474.5 | -4758.9 | 4 (4 + 0) | $R_{A \rightarrow G} = 2.6, R_{A \rightarrow T} = 0.6, R_{C \rightarrow A} = 0.9, R_{C \rightarrow G} = 1.0$ |

**Table 2:** Experimentally informed evolutionary models also provide a superior phylogenetic fit when the tree topology is estimated using the model of Kosiol et al. (2007) rather than that of Goldman and Yang (1994). This table differs from Table 1 in that the phylogenetic fit is to all TEM and SHV sequences using the tree topology in Figure 2B rather than that in Figure 2A.

| model | $\Delta AIC$ | log<br>likelihood | parameters<br>(optimized +<br>empirical) | optimized parameters |
| --- | --- | --- | --- | --- |
| experimental, Equation 2, free $\beta$ | 0.0 | -4019.9 | 5 (5 + 0) | $R_{A \rightarrow G} = 2.2, R_{A \rightarrow T} = 0.5, R_{C \rightarrow A} = 0.8, R_{C \rightarrow G} = 0.8, \beta = 1.6$ |
| experimental, Equation 2, $\beta = 1$ | 48.3 | -4045.1 | 4 (4 + 0) | $R_{A \rightarrow G} = 2.2, R_{A \rightarrow T} = 0.5, R_{C \rightarrow A} = 0.8, R_{C \rightarrow G} = 0.8$ |
| experimental, Equation 3, free $\beta$ | 76.1 | -4058.0 | 5 (5 + 0) | $R_{A \rightarrow G} = 2.1, R_{A \rightarrow T} = 0.5, R_{C \rightarrow A} = 0.8, R_{C \rightarrow G} = 0.8, \beta = 1.3$ |
| experimental, Equation 3, $\beta = 1$ | 85.6 | -4063.7 | 4 (4 + 0) | $R_{A \rightarrow G} = 2.1, R_{A \rightarrow T} = 0.5, R_{C \rightarrow A} = 0.8, R_{C \rightarrow G} = 0.8$ |
| GY94, gamma $\omega$, gamma rates | 398.0 | -4210.9 | 13 (4 + 9) | $\kappa = 2.8, \omega \text{ shape} = 0.4, \text{mean } \omega = 0.6, \text{rate shape} = 1.3$ |
| KOSI07+F, gamma $\omega$, gamma rates | 402.2 | -4162.0 | 64 (4 + 60) | $\kappa = 0.3, \omega \text{ shape} = 0.4, \text{mean } \omega = 4.2, \text{rate shape} = 1.4$ |
| KOSI07+F, gamma $\omega$, one rate | 455.1 | -4189.5 | 63 (3 + 60) | $\kappa = 0.3, \omega \text{ shape} = 0.4, \text{mean } \omega = 4.3$ |
| GY94, gamma $\omega$, one rate | 464.4 | -4245.1 | 12 (3 + 9) | $\kappa = 2.7, \omega \text{ shape} = 0.3, \text{mean } \omega = 0.9$ |
| GY94, one $\omega$, gamma rates | 527.9 | -4276.9 | 12 (3 + 9) | $\kappa = 2.5, \omega = 0.4, \text{rate shape} = 1.2$ |
| KOSI07+F, one $\omega$, gamma rates | 529.8 | -4226.9 | 63 (3 + 60) | $\kappa = 0.4, \omega = 2.0, \text{rate shape} = 1.6$ |
| KOSI07+F, one $\omega$, one rate | 608.3 | -4267.1 | 62 (2 + 60) | $\kappa = 0.4, \omega = 2.0$ |
| GY94, one $\omega$, one rate | 651.9 | -4339.9 | 11 (2 + 9) | $\kappa = 2.3, \omega = 0.4$ |
| randomized, Equation 3, free $\beta$ | 726.3 | -4383.1 | 5 (5 + 0) | $R_{A \rightarrow G} = 2.4, R_{A \rightarrow T} = 0.5, R_{C \rightarrow A} = 0.9, R_{C \rightarrow G} = 0.9, \beta = 0.1$ |
| randomized, Equation 2, free $\beta$ | 737.0 | -4388.4 | 5 (5 + 0) | $R_{A \rightarrow G} = 2.4, R_{A \rightarrow T} = 0.5, R_{C \rightarrow A} = 0.9, R_{C \rightarrow G} = 0.9, \beta = 0.0$ |
| avg. frequencies, Equation 3, free $\beta$ | 823.7 | -4371.8 | 65 (5 + 60) | $R_{A \rightarrow G} = 2.4, R_{A \rightarrow T} = 0.6, R_{C \rightarrow A} = 1.0, R_{C \rightarrow G} = 1.0, \beta = 0.5$ |
| avg. frequencies, Equation 2, free $\beta$ | 844.8 | -4382.3 | 65 (5 + 60) | $R_{A \rightarrow G} = 2.4, R_{A \rightarrow T} = 0.6, R_{C \rightarrow A} = 1.0, R_{C \rightarrow G} = 1.0, \beta = 0.3$ |
| avg. frequencies, Equation 3, $\beta = 1$ | 862.1 | -4392.0 | 64 (4 + 60) | $R_{A \rightarrow G} = 2.3, R_{A \rightarrow T} = 0.6, R_{C \rightarrow A} = 1.1, R_{C \rightarrow G} = 1.0$ |
| avg. frequencies, Equation 2, $\beta = 1$ | 907.1 | -4414.5 | 64 (4 + 60) | $R_{A \rightarrow G} = 2.4, R_{A \rightarrow T} = 0.6, R_{C \rightarrow A} = 1.1, R_{C \rightarrow G} = 1.1$ |
| randomized, Equation 3 | 1265.1 | -4653.5 | 4 (4 + 0) | $R_{A \rightarrow G} = 2.3, R_{A \rightarrow T} = 0.5, R_{C \rightarrow A} = 0.8, R_{C \rightarrow G} = 1.0$ |
| randomized, Equation 2 | 1474.1 | -4758.0 | 4 (4 + 0) | $R_{A \rightarrow G} = 2.5, R_{A \rightarrow T} = 0.5, R_{C \rightarrow A} = 0.8, R_{C \rightarrow G} = 1.1$ |

The experimentally informed models are superior to the non-site-specific models even when the stringency parameter *β* is fixed to one; however, the phylogenetic fit is substantially enhanced by treating *β* as a free parameter (Table 1 and Table 2). In all cases, fitting of the stringency parameter yields values of *β >* 1, indicating that natural evolution is more stringent than the experiments in its preferences for specific amino-acids. This result is consistent with the notion that selection during real evolution selection is more sensitive than the typical laboratory experiment.

To confirm that the superiority of the experimentally informed models is due to the fact that the deep mutational scanning of Firnberg et al. (2014) captures information about the site-specific amino-acid preferences, I tested evolutionary models in which these preferences were randomized among sites or were set to the average frequencies of all amino-acids in the lactamase alignments. In the former case, the *π_r,a_* values are randomly shuffled among sites, where in the latter case *π_r,a_* for all values of *r* is set to the average frequency of amino-acid *a* over the entire lactamase alignment. As can be seen from Table 1 and Table 2, these non-site-specific models perform substantially worse than even the simplest versions of the models of Goldman and Yang (1994) and Kosiol et al. (2007). This result shows that the improved performance of the experimentally informed evolutionary models is overwhelmingly due to incorporation of information on the site-specific amino-acid preferences rather than better modeling of the mutational process.

#### Experimentally informed models are superior for individual TEM and SHV phylogenies

The foregoing results show that experimentally informed models are superior for describing the combined TEM and SHV beta-lactamase phylogeny. Given that the amino-acid preferences were determined by experiments using a TEM-1 parent, it is worth asking whether these preferences accurately describe the evolution of both the TEM and SHV sequences, or whether they more accurately describe the TEM sequences (which are closely related to TEM-1, Figure 2) than the SHV sequences (which only have about 65% protein identity to TEM-1, Figure 2). This question is relevant because the extent to which site-specific amino-acid preferences are conserved during protein evolution remains unclear. For instance, while several experimental studies have suggested that such preferences are largely conserved among moderately diverged homologs (Ashenberg et al., 2013; Serrano et al., 1993), a simulation-based study has argued that preferences shift substantially during protein evolution (Pollock et al., 2012; Pollock and Goldstein, 2014). If the site-specific amino-acid preferences are largely conserved during the divergence of the TEM and SHV sequences, then the experimentally informed models should work well for both these groups – but if the preferences shift rapidly during evolution, then the experimentally informed models should be effective only for the closely related TEM sequences.

To test these competing possibilities, I repeated the analysis in the foregoing section separately for the TEM and SHV clades of the overall phylogenetic tree (the red versus blue clades in Figure 2). This analysis found that the experimentally informed evolutionary models were clearly superior to all alternative models for the SHV as well as the TEM clade (Table 3, Table 4, Table 5, Table 6). In fact, the extent of superiority of the experimentally informed model (as quantified by AIC) was greater for the SHV clade than the TEM clade, despite the fact that the former has fewer sequences. These results suggest that the applicability of the experimentally determined amino-acid preferences extends to beta-lactamase homologs that are substantially diverged from the TEM-1 parent that was the specific subject of the experiments of Firnberg et al. (2014).

**Table 3:** Experimentally informed evolutionary models also provide a superior phylogenetic fit when the analysis is limited only to TEM beta-lactamase sequences. This table differs from Table 1 in that the phylogenetic fit is only to the TEM sequences (the portion of the tree shown in red in Figure 2A.)

| model | ΔAIC | log likelihood | parameters (optimized + empirical) | optimized parameters |
| --- | --- | --- | --- | --- |
| experimental, Equation 2, free $\beta$ | 0.0 | -2374.3 | 5 (5 + 0) | $R_{A \rightarrow G} = 1.9, R_{A \rightarrow T} = 0.4, R_{C \rightarrow A} = 1.1, R_{C \rightarrow G} = 0.5, \beta = 1.5$ |
| experimental, Equation 2, $\beta = 1$ | 23.1 | -2386.8 | 4 (4 + 0) | $R_{A \rightarrow G} = 1.9, R_{A \rightarrow T} = 0.4, R_{C \rightarrow A} = 1.1, R_{C \rightarrow G} = 0.5$ |
| experimental, Equation 3, free $\beta$ | 81.8 | -2415.2 | 5 (5 + 0) | $R_{A \rightarrow G} = 1.8, R_{A \rightarrow T} = 0.4, R_{C \rightarrow A} = 1.1, R_{C \rightarrow G} = 0.5, \beta = 1.2$ |
| experimental, Equation 3, $\beta = 1$ | 83.6 | -2417.1 | 4 (4 + 0) | $R_{A \rightarrow G} = 1.8, R_{A \rightarrow T} = 0.4, R_{C \rightarrow A} = 1.1, R_{C \rightarrow G} = 0.5$ |
| GY94, gamma $\omega$, gamma rates | 252.2 | -2492.4 | 13 (4 + 9) | $\kappa = 3.1, \omega \text{ shape} = 0.3, \text{mean } \omega = 1.3, \text{rate shape} = 0.4$ |
| GY94, one $\omega$, gamma rates | 317.6 | -2526.1 | 12 (3 + 9) | $\kappa = 3.0, \omega = 1.1, \text{rate shape} = 0.3$ |
| GY94, gamma $\omega$, one rate | 318.5 | -2526.5 | 12 (3 + 9) | $\kappa = 3.2, \omega \text{ shape} = 0.2, \text{mean } \omega = 1.6$ |
| KOSI07+F, gamma $\omega$, gamma rates | 326.9 | -2478.7 | 64 (4 + 60) | $\kappa = 0.3, \omega \text{ shape} = 0.3, \text{mean } \omega = 10.4, \text{rate shape} = 0.4$ |
| KOSI07+F, gamma $\omega$, one rate | 394.9 | -2513.8 | 63 (3 + 60) | $\kappa = 0.3, \omega \text{ shape} = 0.2, \text{mean } \omega = 13.7$ |
| KOSI07+F, one $\omega$, gamma rates | 412.0 | -2522.3 | 63 (3 + 60) | $\kappa = 0.3, \omega = 7.9, \text{rate shape} = 0.3$ |
| randomized, Equation 3, free $\beta$ | 465.8 | -2607.2 | 5 (5 + 0) | $R_{A \rightarrow G} = 2.1, R_{A \rightarrow T} = 0.5, R_{C \rightarrow A} = 1.2, R_{C \rightarrow G} = 0.5, \beta = 0.0$ |
| randomized, Equation 2, free $\beta$ | 466.2 | -2607.4 | 5 (5 + 0) | $R_{A \rightarrow G} = 2.1, R_{A \rightarrow T} = 0.5, R_{C \rightarrow A} = 1.2, R_{C \rightarrow G} = 0.5, \beta = 0.0$ |
| GY94, one $\omega$, one rate | 483.3 | -2609.9 | 11 (2 + 9) | $\kappa = 2.9, \omega = 1.1$ |
| KOSI07+F, one $\omega$, one rate | 556.7 | -2595.7 | 62 (2 + 60) | $\kappa = 0.3, \omega = 7.2$ |
| avg. frequencies, Equation 2, free $\beta$ | 574.6 | -2601.6 | 65 (5 + 60) | $R_{A \rightarrow G} = 2.1, R_{A \rightarrow T} = 0.5, R_{C \rightarrow A} = 1.2, R_{C \rightarrow G} = 0.5, \beta = 0.3$ |
| avg. frequencies, Equation 3, free $\beta$ | 577.9 | -2603.2 | 65 (5 + 60) | $R_{A \rightarrow G} = 2.1, R_{A \rightarrow T} = 0.5, R_{C \rightarrow A} = 1.2, R_{C \rightarrow G} = 0.5, \beta = 0.3$ |
| avg. frequencies, Equation 2, $\beta = 1$ | 609.1 | -2619.8 | 64 (4 + 60) | $R_{A \rightarrow G} = 2.1, R_{A \rightarrow T} = 0.5, R_{C \rightarrow A} = 1.3, R_{C \rightarrow G} = 0.6$ |
| avg. frequencies, Equation 3, $\beta = 1$ | 622.7 | -2626.7 | 64 (4 + 60) | $R_{A \rightarrow G} = 2.0, R_{A \rightarrow T} = 0.5, R_{C \rightarrow A} = 1.3, R_{C \rightarrow G} = 0.6$ |
| randomized, Equation 3 | 976.6 | -2863.6 | 4 (4 + 0) | $R_{A \rightarrow G} = 2.0, R_{A \rightarrow T} = 0.5, R_{C \rightarrow A} = 1.1, R_{C \rightarrow G} = 0.5$ |
| randomized, Equation 2 | 1007.8 | -2879.2 | 4 (4 + 0) | $R_{A \rightarrow G} = 2.3, R_{A \rightarrow T} = 0.5, R_{C \rightarrow A} = 1.1, R_{C \rightarrow G} = 0.5$ |

**Table 4:** Experimentally informed evolutionary models also provide a superior phylogenetic fit when the analysis is limited only to SHV beta-lactamase sequences. This table differs from Table 1 in that the phylogenetic fit is only to the SHV sequences (the portion of the tree shown in blue in Figure 2A.)

| model | ΔAIC | log<br>likelihood | parameters<br>(optimized +<br>empirical) | optimized parameters |
| --- | --- | --- | --- | --- |
| experimental, Equation 2, free $\beta$ | 0.0 | -1728.5 | 5 (5 + 0) | $R_{A \rightarrow G} = 4.4, R_{A \rightarrow T} = 1.9, R_{C \rightarrow A} = 0.5, R_{C \rightarrow G} = 1.1, \beta = 2.5$ |
| experimental, Equation 3, free $\beta$ | 34.9 | -1746.0 | 5 (5 + 0) | $R_{A \rightarrow G} = 4.2, R_{A \rightarrow T} = 1.9, R_{C \rightarrow A} = 0.5, R_{C \rightarrow G} = 1.1, \beta = 2.2$ |
| experimental, Equation 2, $\beta = 1$ | 106.2 | -1782.7 | 4 (4 + 0) | $R_{A \rightarrow G} = 4.5, R_{A \rightarrow T} = 1.9, R_{C \rightarrow A} = 0.6, R_{C \rightarrow G} = 1.2$ |
| experimental, Equation 3, $\beta = 1$ | 116.5 | -1787.8 | 4 (4 + 0) | $R_{A \rightarrow G} = 4.4, R_{A \rightarrow T} = 1.9, R_{C \rightarrow A} = 0.6, R_{C \rightarrow G} = 1.2$ |
| KOSI07+F, gamma $\omega$, gamma rates | 489.0 | -1914.0 | 64 (4 + 60) | $\kappa = 0.2, \omega \text{ shape} = 0.5, \text{mean } \omega = 2.9, \text{rate shape} = 0.2$ |
| KOSI07+F, one $\omega$, gamma rates | 499.7 | -1920.4 | 63 (3 + 60) | $\kappa = 0.2, \omega = 1.8, \text{rate shape} = 0.3$ |
| GY94, gamma $\omega$, gamma rates | 505.8 | -1973.4 | 13 (4 + 9) | $\kappa = 3.6, \omega \text{ shape} = 0.6, \text{mean } \omega = 0.5, \text{rate shape} = 0.2$ |
| GY94, one $\omega$, gamma rates | 514.0 | -1978.5 | 12 (3 + 9) | $\kappa = 3.4, \omega = 0.3, \text{rate shape} = 0.2$ |
| KOSI07+F, gamma $\omega$, one rate | 555.7 | -1948.4 | 63 (3 + 60) | $\kappa = 0.3, \omega \text{ shape} = 0.3, \text{mean } \omega = 2.0$ |
| KOSI07+F, one $\omega$, one rate | 573.6 | -1958.3 | 62 (2 + 60) | $\kappa = 0.3, \omega = 1.8$ |
| GY94, gamma $\omega$, one rate | 581.5 | -2012.3 | 12 (3 + 9) | $\kappa = 3.4, \omega \text{ shape} = 0.3, \text{mean } \omega = 0.4$ |
| randomized, Equation 3, free $\beta$ | 601.7 | -2029.4 | 5 (5 + 0) | $R_{A \rightarrow G} = 5.0, R_{A \rightarrow T} = 1.8, R_{C \rightarrow A} = 0.6, R_{C \rightarrow G} = 1.4, \beta = 0.1$ |
| GY94, one $\omega$, one rate | 602.6 | -2023.8 | 11 (2 + 9) | $\kappa = 3.4, \omega = 0.3$ |
| randomized, Equation 2, free $\beta$ | 602.7 | -2029.9 | 5 (5 + 0) | $R_{A \rightarrow G} = 5.1, R_{A \rightarrow T} = 1.8, R_{C \rightarrow A} = 0.6, R_{C \rightarrow G} = 1.5, \beta = 0.0$ |
| avg. frequencies, Equation 3, free $\beta$ | 711.5 | -2024.3 | 65 (5 + 60) | $R_{A \rightarrow G} = 5.0, R_{A \rightarrow T} = 2.0, R_{C \rightarrow A} = 0.7, R_{C \rightarrow G} = 1.5, \beta = 0.3$ |
| avg. frequencies, Equation 2, free $\beta$ | 715.7 | -2026.4 | 65 (5 + 60) | $R_{A \rightarrow G} = 5.1, R_{A \rightarrow T} = 1.9, R_{C \rightarrow A} = 0.7, R_{C \rightarrow G} = 1.5, \beta = 0.3$ |
| avg. frequencies, Equation 3, $\beta = 1$ | 749.8 | -2044.5 | 64 (4 + 60) | $R_{A \rightarrow G} = 4.9, R_{A \rightarrow T} = 2.1, R_{C \rightarrow A} = 0.8, R_{C \rightarrow G} = 1.6$ |
| avg. frequencies, Equation 2, $\beta = 1$ | 758.8 | -2048.9 | 64 (4 + 60) | $R_{A \rightarrow G} = 5.2, R_{A \rightarrow T} = 2.1, R_{C \rightarrow A} = 0.8, R_{C \rightarrow G} = 1.7$ |
| randomized, Equation 3 | 1047.0 | -2253.1 | 4 (4 + 0) | $R_{A \rightarrow G} = 4.8, R_{A \rightarrow T} = 1.8, R_{C \rightarrow A} = 0.6, R_{C \rightarrow G} = 1.4$ |
| randomized, Equation 2 | 1071.2 | -2265.2 | 4 (4 + 0) | $R_{A \rightarrow G} = 5.3, R_{A \rightarrow T} = 1.7, R_{C \rightarrow A} = 0.6, R_{C \rightarrow G} = 1.5$ |

**Table 5:**
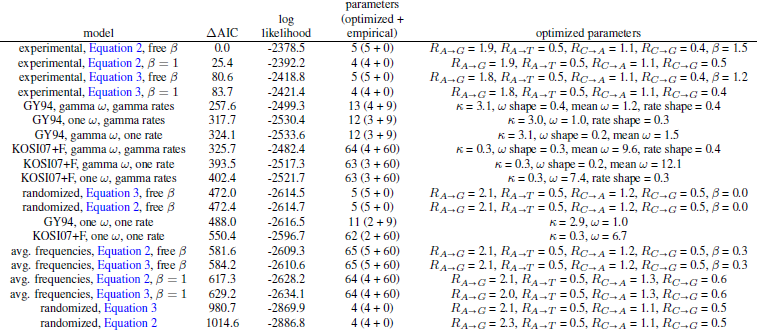
Experimentally informed evolutionary models also provide a superior phylogenetic fit to the TEM beta-lactamases when the tree topology is estimated using the model of Kosiol et al. (2007) rather than that of Goldman and Yang (1994). This table differs from Table 3 in that the phylogenetic fit is to the TEM sequences using the red portion of tree topology in Figure 2B rather than the red portion of the tree topology in Figure 2A.

**Table 6:**
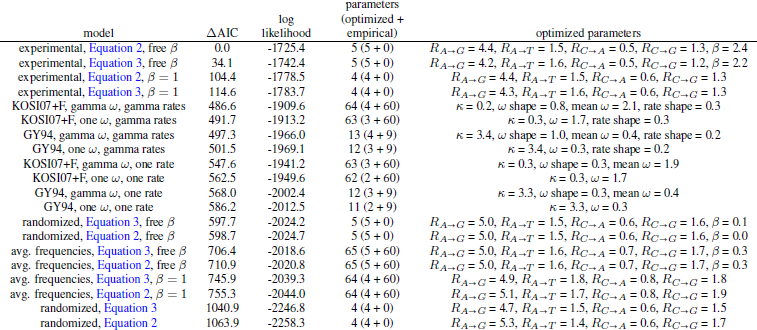
Experimentally informed evolutionary models also provide a superior phylogenetic fit to the SHV beta-lactamases when the tree topology is estimated using the model of Kosiol et al. (2007) rather than that of Goldman and Yang (1994). This table differs from Table 4 in that the phylogenetic fit is to the SHV sequences using the blue portion of tree topology in Figure 2B rather than the blue portion of the tree topology in Figure 2A.

#### The stringency parameter ***β*** generally improves experimentally informed models

The results in the foregoing sections show that use of the stringency parameter *β* improves the phylogenetic fit of the experimentally informed models of lactamase evolution. Previous work (Bloom, 2014) has shown that an experimentally informed evolutionary model without a stringency parameter (*β* = 1) improves the phylogenetic fit to influenza nucleoprotein sequences relative to non-site-specific models. To test whether the fitting of a stringency parameter further improves the experimentally informed evolutionary model of nucleoprotein, I repeated the analysis of Bloom (2014) but also included a model variant in which *β* was fit by maximum likelihood.

Table 7 shows that fitting the stringency parameter substantially improves the phylogenetic fit of the experimentally informed model of nucleoprotein evolution. This table also shows that as with the lactamase models described above, the fitted value of *β* was greater than one. Overall, this result suggests that inclusion of a stringency parameter generally improves the phylogenetic fit of experimentally informed evolutionary models. Presumably this improvement arises from the fact that deep mutational scanning experiments are generally less stringent than real evolution, and so values of *β >* 1 help scale the experimentally determined site-specific amino-acid preferences to a stringency more reflective of those that constrain real evolution.

**Table 7:** Fitting of the stringency parameter *β* also improves the phylogenetic fit of an experimentally informed evolutionary model for influenza nucleoprotein. The data in this table were generated by exactly repeating the analysis in the sixth table of Bloom (2014) except including the additional models listed here, which include a model with a stringency parameter. With the exception of the stringency parameter, the experimentally informed evolutionary model used here is derived entirely from experimental measurements, since both the mutation rates and amino-acid preferences were measured in Bloom (2014). Only the model using fixation probabilities calculated from Equation 3 are reported, since Bloom (2014) shows that this is the best model for influenza nucleoprotein. The data and source code used to generate this table are available via http://jbloom.github.io/phyloExpCM/example_2014Analysis_Influenza_NP_Human_1918_Descended_withbeta.html.

| model | $\Delta AIC$ | log<br>likelihood | parameters<br>(optimized +<br>empirical) | optimized parameters |
| --- | --- | --- | --- | --- |
| experimental, Equation 3, free $\beta$ | 0.0 | -12144.2 | 1 (1 + 0) | $\beta = 1.7$ |
| experimental, Equation 3, $\beta = 1$ | 391.8 | -12341.1 | 0 (0 + 0) | none |
| GY94, gamma $\omega$, gamma rates | 1453.0 | -12858.7 | 13 (4 + 9) | $\kappa = 5.9$, $\omega$ shape = 0.2, mean $\omega = 0.2$, rate shape = 2.5 |
| GY94, gamma $\omega$, one rate | 1616.0 | -12941.2 | 12 (3 + 9) | $\kappa = 5.6$, $\omega$ shape = 0.2, mean $\omega = 0.2$ |
| KOSI07+F, gamma $\omega$, gamma rates | 1845.6 | -13004.0 | 64 (4 + 60) | $\kappa = 0.2$, $\omega$ shape = 0.2, mean $\omega = 0.9$, rate shape = 1.8 |
| GY94, one $\omega$, gamma rates | 1884.1 | -13075.3 | 12 (3 + 9) | $\kappa = 5.9$, $\omega = 0.1$, rate shape = 2.0 |
| GY94, one $\omega$, one rate | 2153.2 | -13210.8 | 11 (2 + 9) | $\kappa = 5.5$, $\omega = 0.1$ |
| KOSI07+F, gamma $\omega$, one rate | 2153.6 | -13159.0 | 63 (3 + 60) | $\kappa = 0.2$, $\omega$ shape = 0.2, mean $\omega = 1.2$ |
| KOSI07+F, one $\omega$, gamma rates | 2227.3 | -13195.9 | 63 (3 + 60) | $\kappa = 0.2$, $\omega = 0.7$, rate shape = 1.6 |
| KOSI07+F, one $\omega$, one rate | 2650.1 | -13408.3 | 62 (2 + 60) | $\kappa = 0.2$, $\omega = 0.7$ |
| avg. frequencies, Equation 3, free $\beta$ | 3736.2 | -13952.3 | 61 (1 + 60) | $\beta = 1.2$ |
| avg. frequencies, Equation 3, $\beta = 1$ | 3742.2 | -13956.3 | 60 (0 + 60) | none |

#### Experimentally informed models are slightly better for many sites

The results described above and in Bloom (2014) demonstrate that experimentally informed evolutionary models improve phylogenetic relative to non-site-specific models when analyzing the entire gene sequences of lactamase or nucleoprotein. It is interesting to investigate *which* sites are described more accurately by the experimentally informed models. This can be done by examining the differences in per-site likelihoods between models after fixing the branch lengths and model parameters to their maximum likelihood values for the entire gene.

Figure 3 compares the per-site likelihoods for the best experimentally informed evolutionary model for lactamase and for nucleoprotein relative to the best non-site-specific model for each of these genes (using the models from Table 1 and Table 7). For both lactamase and nucleoprotein, the experimentally informed models lead to small improvements for many sites. Overall, 72% (207 of 286) lactamase sites are described better by the experimentally informed model (Supplementary file 3), and 82% (407 of 498) of nucleoprotein sites are described better by the experimentally informed model (Supplementary file 4). There appears to be a slight trend for improvements due to the experimentally informed models to be most common for buried sites, but the experimentally informed models lead to small improvements for sites spanning a range of solvent accessibilities and secondary structures.

**Figure 3:**
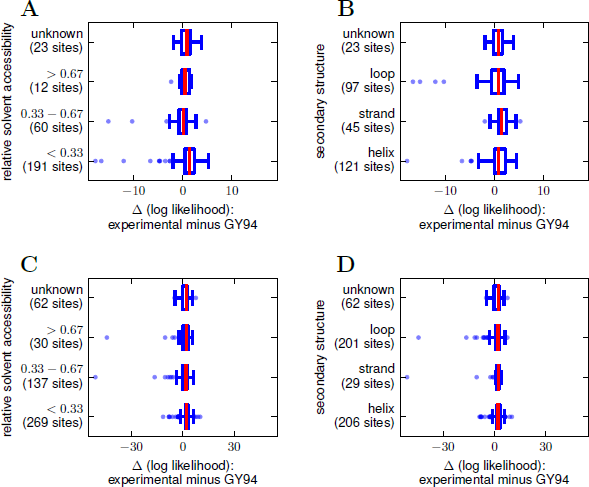
Comparison of likelihoods on a per-site basis between the best experimentally informed site-specific evolutionary model and the best conventional non-site-specific model. The experimentally informed models are slightly better (positive Δ (log likelihood)) for most sites, but far worse for a handful of sites. **(A), (B)** The best experimentally informed lactamase model in Table 1 versus the best GY94 variant in Table 1. The experimentally informed model has a higher log likelihood for 72% of lactamase sites. **(C), (D)** The best experimentally informed nucleo-protein model in Table 7 versus the best GY94 variant in Table 7. The experimentally informed model has a higher log likelihood for 82% of sites. For both genes, the per-site likelihoods were computed after fixing the model parameters and branch lengths to their maximum-likelihood values for the entire gene. Sites are classified in terms of their relative solvent accessibility or secondary structure as computed using DSSP (Kabsch and Sander, 1983; Joosten et al., 2011) from PDB structures 1XPB (Fonzé et al., 1995) or 2IQH (Ye et al., 2006), normalizing solvent accessibilities to the values provided by Tien et al. (2013). The per-residue numerical data are in Supplementary file 3 and Supplementary file 4. The code and data used to create this figure are provided via http://jbloom.github.io/phyloExpCM/example_2014Analysis_lactamase.html and http://jbloom.github.io/phyloExpCM/example_2014Analysis_Influenza_NP_Human_1918_Descended_withbeta.html.

Interestingly, for both genes there are also a few sites for which the experimentally informed models are *far worse* than the non-site-specific models. Presumably these sites are modeled poorly because the preferences determined by the deep mutational scanning experiments do not accurately capture the real preferences of natural evolution. Such discrepancies could arise from either a failure of the deep mutational scanning experiments to fully reflect natural selection pressures, or from epistatic interactions that strongly shift the site preferences during natural evolution away from those measured for the parent gene by the experiments. I was unable to observe any obvious features of the specific sites that were described poorly by the experimentally informed models (Supplementary file 3, Supplementary file 4). It therefore remains an open question why certain sites are described very poorly by the experimentally informed models even though the vast majority of sites are described better by these models.

#### Comparison of different methods for computing fixation probabilities

In the foregoing analyses, two different mathematical relationships were used to mathematically relate the experimentally determined amino-acid preferences to the substitution probabilities in the evolutionary models. One relationship (Equation 2) is based on a true population-genetics derivation by Halpern and Bruno (1998) under the assumption that the preferences are reflective of selection coefficients for amino acids at specific sites (an assumption that would only be strictly true only in the unlikely case that individual sites in a gene contribute independently to fitness). The other relationship (Equation 3) is a more *ad hoc* one that I suggested in previous work (Bloom, 2014) on the grounds that the amino-acid preferences might be best envisioned not as selection coefficients, but rather as measurements of the *fraction* of genetic backgrounds that tolerate a specific mutation, as would be implied by the evolutionary dynamics described in Bloom et al. (2007). Although both relationships share the qualitative feature that mutations to higher-preference amino acids are favored over mutations to lower-preference ones, they differ in their quantitative details. In previous work on influenza nucleoprotein (Bloom, 2014), I reported that the relationship in Equation 3 outperformed the one in Equation 2 derived by Halpern and Bruno (1998).

In contrast, for the beta-lactamase sequences studied here, the relationship of Halpern and Bruno (1998) outperforms the one in Equation 3 (Table 1, Table 2, Table 3, Table 4, Table 5, Table 6). The reason for and relevance of these discordant results remains unclear. There are almost certainly differences in the evolutionary conditions (population size, degree of polymorphism, etc) for influenza nucleoprotein and beta-lactamase that influence the relationship between selection coefficients and fixation probabilities. In addition, there are substantial differences between the experiments of Firnberg et al. (2014) on beta-lactamase and my previous work on nucleoprotein– in particular, Firnberg et al. (2014) examine the effects of single mutations to the parental gene, whereas the nucleoprotein experiments examined the average effects of individual mutations in variants that often contained multiple mutations. Finally, the experimental measurements are imperfect – for nucleoprotein, the preferences determined by independent biological replicates of the experiments only had a Pearson correlation coefficient of 0.79; Firnberg et al. (2014) do not provide data on the consistency of their measurements across biological replicates, but it seems safe to assume that their experiments are also imperfect. Therefore, further work is probably needed to determine if any meaning can be ascribed to the differences in fit for Equation 2 versus Equation 3, as well as to identify the optimal mathematical relationship for connecting experimentally measured amino-acid preferences to substitution probabilities in evolutionary models. However, both the results presented here and in Bloom (2014) strongly suggest that using any reasonable mathematical relationship to inform evolutionary models with experimentally determined amino-acid preferences is sufficient to lead to dramatic improvements in phylogenetic fit.

It is also interesting to speculate on the precise meaning of the stringency parameter *β*. According to population-genetics theory, the strength of selection increases with effective population size. It is therefore tempting to interpret *β* as reflecting differences in the effective population size in the deep mutational scanning experiments relative to natural evolution. Indeed, one could imagine even attempting to use the inferred value of *β* to indirectly quantify effective population size. However, it is important to temper this tantalizing possibility with the reminder that the fixation probabilities over the entire phylogeny can be related to the fitness effects of specific mutations only under the unrealistic assumption that the fitness contributions of mutations at different sites are entirely independent, since these probabilities reflect the effect of a mutation averaged over all genetic backgrounds in the phylogeny. So while the fact that using *β >* 1 consistently improves phylogenetic does indicate that natural evolution is more stringent in its site-specific amino-acid preferences than the experiments, the exact population-genetics interpretation of *β* is unclear.

## Discussion

I have shown that an evolutionary model informed by experimentally determined site-specific amino-acid preferences fits beta-lactamase phylogenies better than a variety of existing models that do not utilize site-specific information. When considered in combination with prior work demonstrating that an experimentally determined evolutionary model dramatically improves phylogenetic fit for influenza nucleoprotein (Bloom, 2014), these results suggest that experimentally informed models are generally superior to non-site-specific models for phylogenetic analyses of protein-coding genes. The explanation for this superiority is obvious: proteins have strong preferences for certain amino acids at specific sites (Ashenberg et al., 2013; Halpern and Bruno, 1998) which are ignored by non-site-specific models. The use of experimentally measured site-specific amino-acid preferences improves evolutionary models by informing them about the complex selection that shapes actual sequence evolution. Interestingly, inclusion of a parameter that allows the stringency of site-specific amino-acid preferences during natural evolution to exceed those measured by experiments further improves phylogenetic fit, suggesting that deep mutational scanning experiments remain less sensitive than actual natural selection.

I have not compared the experimentally informed evolutionary models to more recent site-specific models that infer aspects of the substitution process from the sequence data itself (Lartillot and Philippe, 2004; Rodrigue et al., 2010; Le et al., 2008; Wu et al., 2013; Wang et al., 2008). Such a comparison is desirable since these site-specific models will almost certainly compare more favorably to the experimentally informed models used here. Unfortunately, such a comparison is sufficiently technically challenging to be beyond the scope of the current work, since it requires comparing between maximum-likelihood and Bayesian approaches. In any case, the experimentally informed evolutionary models described here should not be viewed purely as competitors for existing site-specific substitution models. Rather, one can imagine future approaches that integrate the results of a deep mutational scanning experiment with additional site-specific details inferred from natural sequence data to create even more nuanced evolutionary models.

An advantage of experimentally informed evolutionary models is that they naturally lend themselves to interpretation in terms of quantities that can be directly related to both specific biochemical measurements and the underlying processes of mutation and selection. This stands in contrast to most existing models, which are phrased in terms of heuristic parameters (such as codon “equilibrium frequencies”) that reflect the combined action of mutation and selection and are not accessible to direct experimental measurement. Experimentally informed evolutionary models therefore have the potential to facilitate connections between the phylogenetic substitution processes and the underlying biochemistry and population genetics of gene evolution (Thorne et al., 2007; Halpern and Bruno, 1998).

The major drawback of experimentally informed models is their more limited scope. Most existing codon-based evolutionary models can be applied to any gene (Goldman and Yang, 1994; Muse and Gaut, 1994; Kosiol et al., 2007) – but experimentally informed models require experimental data for the gene in question. However, this requirement may not be as crippling as it initially appears. The first experimentally determined evolutionary model for influenza nucleoprotein required direct measurement of both the site-specific amino-acid preferences and the underlying mutation rates (Bloom, 2014). However, the model presented here only requires measurement of the amino-acid preferences, as the mutation rates are treated as free parameters. Rapid advances in the experimental technique of deep mutational scanning are making such data available for an increasing number of proteins (Fowler et al., 2010; Roscoe et al., 2013; Starita et al., 2013; Melamed et al., 2013; Traxlmayr et al., 2012; McLaughlin Jr et al., 2012; Firnberg et al., 2014; Bloom, 2014).

In this respect, it is encouraging that the site-specific amino-acid preferences determined experimentally for TEM-1 improve phylogenetic fit to substantially diverged (35% protein sequence divergence) SHV beta-lactamases as well as highly similar TEM beta-lactamases. As discussed in the Introduction, there are two major limitations to most existing evolutionary models: they treat sites identically, and they treat sites independently. Experimentally informed evolutionary models of the type described here have the potential to completely remedy the first limitation as deep mutational scanning defines site-specific selection with increasing precision. However, such models still treat sites independently – and this limitation will never be completely overcome by experiments, since the unforgiving math of combinatorics means that no experiment can examine all arbitrary combinations of mutations (for example, TEM-1 has only 5453 single amino-acid mutants, but it has 14815801 double mutants, 26742520805 triple mutants, and over 10^13^ quadruple mutants). The utility of experimentally informed evolutionary models therefore depends on the extent to which site-specific amino-acid preferences measured for one protein can be extrapolated to other homologs – in other words, are sites sufficiently independent that the preferences at a given position remain similar after mutations at other positions? This question remains a topic of active debate, with experimental studies suggesting that site-specific preferences are largely conserved among closely and moderately related homologs (Ashenberg et al., 2013; Serrano et al., 1993), but some computational studies emphasizing substantial shifts in preferences during evolution (Pollock et al., 2012; Pollock and Goldstein, 2014). The fact that the TEM-1 experimental data informs a model that accurately describes the substantially diverged SHV homologs suggests reasonable conservation of site-specific amino-acid preferences among beta-lactamase homologs.

This apparent conservation of site-specific amino-acid preferences suggests that the phylogenetic utility of experimentally informed evolutionary models may extend well beyond the immediate proteins that were experimentally characterized. This type of experimental generalization would have precedent: only a tiny fraction of proteins have been crystallized, but because structure is largely conserved during protein evolution, it is frequently possible to use a structure determined for one protein to draw insights about a range of related homologs (Lesk and Chothia, 1980; Sander and Schneider, 1991). It seems plausible that the conservation of site-specific amino-acid preferences could similarly enable deep mutational scanning to provide the experimental data to inform evolutionary models of sufficient scope to improve the accuracy and interpretability of phylogenetic analyses for a substantial number of proteins of interest.

## Methods

### Availability of computer code and data

The phylogenetic analyses were performed using the software package *phyloExpCM* (**phylo**genetic analyses with **exp**erimental **c**odon **m**odels, https://github.com/jbloom/phyloExpCM), which primarily serves as an interface to run *HYPHY* (Pond et al., 2005). Input data, computer code, and a description sufficient to enable replication of all analyses reported in this paper are available via http://jbloom.github.io/phyloExpCM/example_2014Analysis_lactamase.html and http://jbloom.github.io/phyloExpCM/example_2014Analysis_Influenza_NP_Human_1918_Descended_withbeta.html.

### Equilibrium frequencies and reversibility

Here I show that the evolutionary model defined by Equation 1 is reversible (satisfies detailed balance), and has *p_r,x_* defined by Equation 9 as its equilibrium frequency.

First, note that the fixation probabilities *F_r,xy_* defined by both Equation 2 and Equation 3 satisfy reversibility with respect to the stringency-adjusted amino-acid preferences 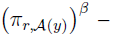 namely that

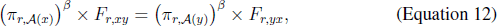

as can be verified by direct substitution. This relationship means that if all codon interchanges were equally likely (all *Q_xy_* values are equal), then the equilibrium frequency *p_r,x_* of codon *x* would simply be proportional to the stringency-adjusted preference 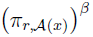 for the encoded amino acid.

However, in practice all codon interchanges are not equally likely, so the actual equilibrium frequencies *p_r,x_* will also depend on the mutation rate parameters *R_m→n_* listed in Equation 8. This dependence is given by the *q_x_* terms in Equation 9. These *q_x_* terms can be thought of as the expected equilibrium frequencies of the codons in a hypothetical situation in which there is no selection and all 64 codons are equally fit (all *F_r,xy_* values are equal). In other words, the *q_x_* terms define the stationary state of the reversible stochastic process defined by the mutation rates *Q_xy_*. In order to show *q_x_* given Equation 10 defines this stationary state, it is necessary to show that

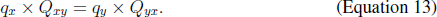

There are up to 12 possible types of single-nucleotide mutations that can be made to a codon *x* to create a different codon *y* (three possible mutations to each of the four possible nucleotides); however only four of these types of mutations require independent verification o f E quation 13. Specifically, the possible types of mutations that require independent verification of Equation 13 are when *x* differs from *y* by a mutation of *A →T,* of *C →G*, of *A →C*, or of *A →G*. The other eight types of mutations do not have to be verified because they are equivalent to one of these first four cases: *C → A* is equivalent to *A* → *C* by symmetry (i.e. interchange of the labels of codons *x* and *y*), *G →A* is equivalent to *A* → *G* by symmetry, *T* → *A* is equivalent to *A* → *T* by symmetry, *G* → *C* is equivalent to *C* → *G* by symmetry, *T* → *G* is equivalent to *A* → *C* because of Equation 5, *T* → *C* is equivalent to *A* → *G* because of Equation 5, *G* → *T* is equivalent to *C* → *A* by Equation 5 and then to *A* → *C* by symmetry, and *C* → *T* is equivalent to *G* → *A* by Equation 5 and then *A* → *G* by symmetry. So below I verify Equation 13 for the four independent types of mutations, full verifying Equation 13.

The first case is where *x* differs from *y* by a mutation of *A* → *T,* such that *Q _xy_* = *R _A_*→*_T_* and *Q_yx_* = *R_T_*_→_*_A_*. In this case, we have

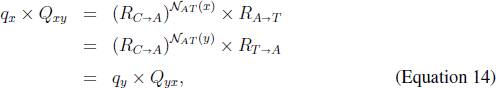

where the first l ine substitutes the definitions of Equation 10 and Equation 4, the second line follows from Equation 5 and the fact that 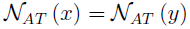 if *x* and *y* differ only by an *A* →*T* mutation, and the final line again substitutes the definitions of Equation 10 and Equation 4.

The second case is where *x* differs from *y* by a mutation of *C → G*, such that *Q_xy_* = *R_C_*_→_*_G_* and *Q_yx_* = *R_G_*_→_*_C_*. In this case, we have

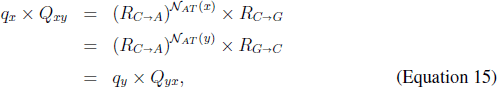

where the justifications for the three lines are identical for those used for Equation 14.

The third case is where *x* differs from *y* by a mutation of *A* _→_ *C*, such that *Q_xy_* = *R_A_*_→_*_C_* and *Q_yx_* = *R_C_*_→_*_A_*. In this case, we have

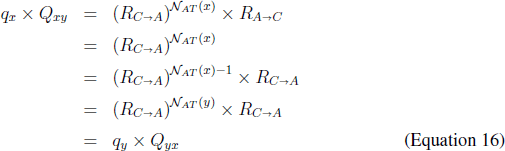

where the first line substitutes the definitions of Equation 10 and Equation 4, the second line uses Equation 7, the third line is simple algebra, the fourth line follows from the fact that 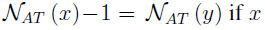 if *x* can be converted to *y* by an *A → C* mutation, and the final line again substitutes the definitions of Equation 10 and Equation 4.

The fourth and final case is where *x* differs from *y* by a mutation of *A → G*, such that *Q_xy_* = *R_A_→_G_* and *Q_y→x_* = *R_G→A_*. In this case, we have

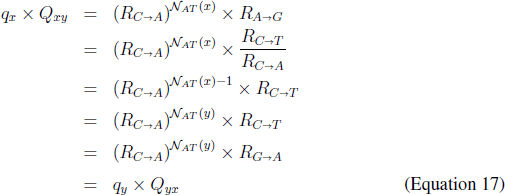

where the first line substitutes the definitions of Equation 10 and Equation 4, the second line uses Equation 6 and Equation 7, the third line is simple algebra, the fourth line follows from the fact that 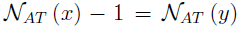 if *x* can be converted to *y* by an *A → C* mutation, the fifth line follows from Equation 5, and the final line again substitutes the definitions of Equation 10 and Equation 4.

Taken together, Equation 14, Equation 15, Equation 16, and Equation 17 establish that Equation 13 holds for all possible independent types of mutations.

Finally, to show that the overall evolutionary model in Equation 1 is reversible and has *p_r,x_* defined by Equation 9 as its equilibrium frequency, it is necessary to show that *p_r,x_ × P_r,xy_* = *p_r,y_ × P_r,yx_*. This follows trivially from Equation 12 and Equation 13:

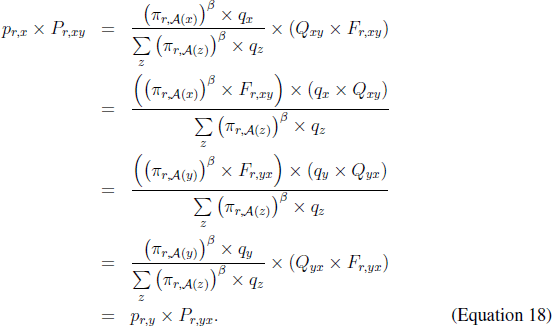

The fact that *P_r,xy_* defines a reversible Markov process with stationary state *p_r,x_* means that it is possible to define a symmetric matrix **S_r_** such that

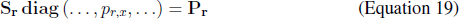

where **diag** (…*, p_r,x_,*…) is the diagonal matrix with *p_r,x_* along its diagonal. Noting 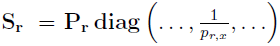, we have

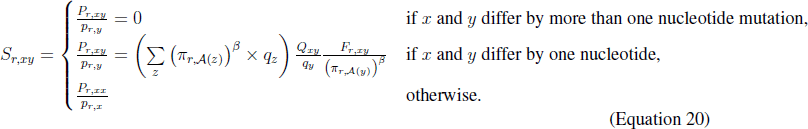

This matrix is symmetric since *S_r,xy_* = *S_r,yx_* as can be verified from the fact that 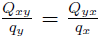 and 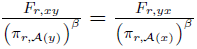 as is guaranteed by Equation 12 and Equation 13.

## Acknowledgments

Thanks to T. Bedford and J. Felsenstein for helpful discussions, and the editor and two anonymous reviewers for exceptionally helpful comments. This work was supported by the National Institute of General Medical Sciences of the National Institutes of Health (grant number R01 GM102198).

---

**Supplementary file 1:** This text file contains the amino-acid preferences displayed graphically in Figure 1. In this file, the amino acids are numbered sequentially starting at one with the N-terminal methionine, rather than using numbering scheme of Ambler et al. (1991) that is employed in Figure 1.

**Supplementary file 2:** This FASTA file contains the alignment of TEM and SHV beta-lactamase sequences used to create the phylogenetic trees in Figure 2.

**Supplementary file 3:** This text file shows the per-site likelihoods for the lactamase phylogeny and their differences between the best experimentally informed evolutionary model and the best variant of the Goldman and Yang model listed in Table 1. Sites are numbered sequentially beginning with 1 at the N-terminal methionine. This is the file *sitelikelihoods.txt* described at http://jbloom.github.io/phyloExpCM/example_2014Analysis_lactamase.html.

**Supplementary file 4:** This text file shows the per-site likelihoods for the nucleoprotein phylogeny and their differences between the best experimentally informed evolutionary model and the best variant of the Goldman and Yang model listed in Table 7. Sites are numbered sequentially beginning with 1 at the N-terminal methionine. This is the file *sitelikelihoods.txt* described at http://jbloom.github.io/phyloExpCM/example_2014Analysis_Influenza_NP_Human_1918_Descended_withbeta.html.

